# Functional characterization of pathway inhibitors for the ubiquitin-proteasome system (UPS) as tool compounds for CRBN and VHL-mediated targeted protein degradation

**DOI:** 10.1101/2024.06.12.598610

**Authors:** Martin P. Schwalm, Amelie Menge, Lewis Elson, Francesco A. Greco, Matthew B. Robers, Susanne Müller, Stefan Knapp

## Abstract

Small molecule degraders such as PROteolysis TArgeting Chimeras (PROTACs) or molecular glues are new modalities for drug development and important tools for target validation. Both modalities recruit an E3 ubiquitin ligase to a protein of interest (POI) either via two independent, but linked ligands (PROTACs) or through binding of a small molecule that alters the E3 binding surface to recruit a neo-substrate (molecular glues). If optimized appropriately, both modalities result in the degradation of the POI. Due to the complexity of the induced multistep degradation process, controls for degrader evaluation are critical and they are commonly used in the literature. However, comparative studies and evaluation of cellular potencies of these control compounds and their appropriate uses have not been published so far. Additionally, the high diversity of mechanisms requires diverse small molecule controls to ensure appropriate inhibition of the investigated system while keeping potential cellular toxicity and unintended effects on cellular pathways as low as possible. Here, we scrutinized a diverse set of ubiquitin pathway inhibitors and evaluated their potency and utility within the CRBN and VHL mediated POI degradation pathway. We used the HiBiT system to measure the levels of target rescue after treatment with control compounds. Additionally, cell health was investigated using a multiplex high content assay. This assay panel allows us to determine non-toxic effective concentrations for control experiments and to perform rescue experiments in the absence of cellular toxicity, which has a profound effect on target degradation by ubiquitin-dependent and -independent pathways.

## Introduction

Targeted protein degradation (TPD) has evolved as a new modality in drug development. Two major compound classes have emerged: I) molecular glues which alter the binding surface of an E3 substrate receptor domain (neo-surface) resulting in a change in substrate selectivity and the recruitment and degradation of neo-substrates; and II) PROteolysis TArgeting Chimeras (PROTACs), which are bivalent (chimeric) compounds binding to a protein of interest (POI) and an E3 ligase ligand, connected by a linker. Compared to molecular glues, PROTACs have a higher molecular weight^1^. The PROTAC dependent recruitment of the E3 ligase to the POI results in a ternary complex that drives POI ubiquitination and subsequent degradation^2,3^. During the past years, many active glues and PROTAC degraders have been developed. However, rational development and optimization of these tools remains challenging due to the multistep nature of ubiquitin-proteasomal system (UPS) mediated degradation of a POI. For efficient target degradation via the UPS, the small molecule-mediated poly-ubiquitin attachment onto the POI requires three molecular events (Figure 1). I) Mono-ubiquitin is activated by E1 enzymes; II) Activated ubiquitin is transferred from the E1 onto an E2; III) The E2-linked ubiquitin then binds to an E3 ligase, followed by ubiquitin transfer either directly onto the target or onto the target after forming an E3-ubiquitin intermediate^4,5^. Additionally, E3 ligases from the cullin-RING family require activation, which is mediated by conjugation of the ubiquitin-like modifier NEDD8 (Figure 1)^6^.

**Figure 1:**
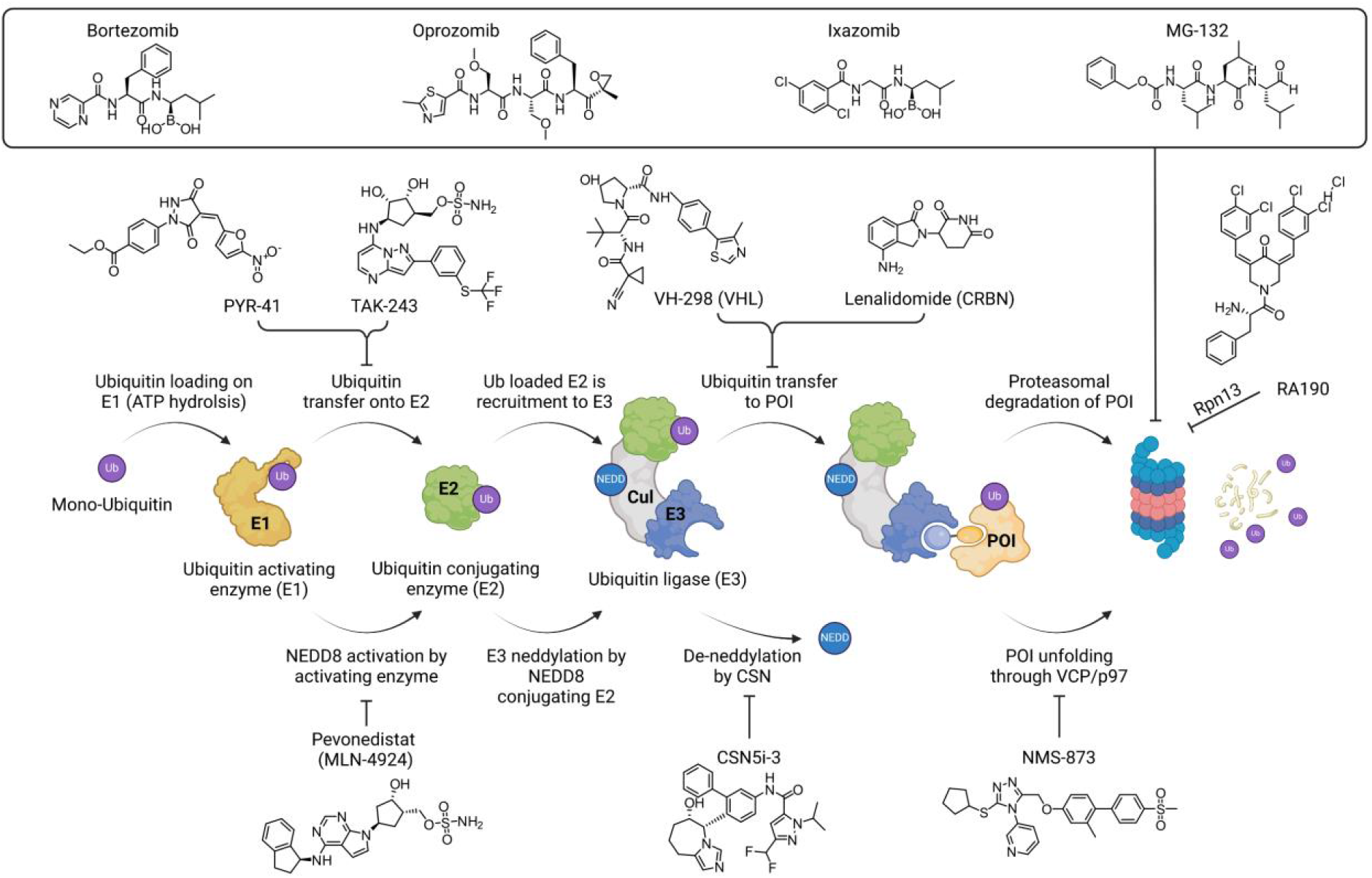
Schematic overview of the small molecule (PROTAC) mediated proteasomal degradation cascade and inhibitors commonly used for the different steps of the cascade.

Generally, changes in protein abundance can be caused by a variety of cellular processes, including modulation of gene transcription, RNA processing and translation as well as physiological protein turnover. Importantly, these processes can be affected by signalling events, changes in the cell cycle and, most importantly, the induction of apoptosis and other cell death mechanisms^7^. It is therefore essential to investigate the molecular mechanism of PROTACs, as they often contain potent inhibitors of essential signalling molecules as warhead structures, that influence different cell states. Especially for PROTACs that are not highly optimized, the cellular on-target inhibition mechanism of the POI may not be negligible, especially when high PROTAC concentrations are used. The protein degradation field has therefore developed several strategies and quality criteria^8^ to control for potentially misleading data, ensuring that the observed phenotypic effect is dependent on the degradation of the POI: Inactive variants of the commonly used E3 ligands CRBN (cereblon) and VHL (von Hippel Lindau) have been developed^9,10^. By incorporating these moieties for PROTAC design instead of active E3 ligands, PROTAC mediated POI degradation is prevented and observed effects, such as POI degradation are instead a consequence due to unrelated mechanisms, such as general toxicity or inhibition of the POI. In addition, proteasome inhibitors such as MG-132 or bortezomib are often used to demonstrate that POI degradation is indeed proteasome-dependent^11^. Less commonly, neddylation inhibitors such as MLN-4924 have been used, demonstrating that the degradation process depends on a cullin E3 systems (CRL)^12,13^. Overall, many chemical tools have been developed targeting essential steps of the UPS. However, these reagents are infrequently used as their utility dissecting the UPS pathway in PROTAC development has not been fully established.

Motivated by the lack of characterisation data on established UPS pathway inhibitors for PROTAC development, we used two well-characterized PROTACs, both potently degrading bromodomain-containing protein 4 (BRD4). The first one, MZ1, uses the pan-BET (BRD2, BRD3, BRD4, BRDT) inhibitor JQ1^14^ linked by a polyethylenglycol (PEG) linker to an established VHL ligand^15^. As second PROTAC dBET6 was used, which is also a JQ1 based PROTAC, but is connected to a CRBN E3 ligand via an aliphatic linker^16^. BRD4 is a highly degradable protein often used in the development of proof-of principle PROTACs. As the target is well studied in the context of TPD, we also used this system as a model in this study. To monitor BRD4 degradation, we made use of an endogenously BRD4-HiBiT tagged cell line (HEK293T^BRD4-HiBiT^)^17^. Twelve inhibitors interfering with different steps of the UPS pathway were evaluated. PYR-41^18^ and TAK-243^19^, potent E1 inhibitors, were used to demonstrate that the degradation process requires a ubiquitin transfer to the E2 enzyme, proving overall ubiquitin dependency. Using the NEDD8-activating enzyme (NAE) inhibitor MLN-4924^20^ which abrogates the NEDD8 conjugation to CRLs, provides evidence that PROTAC mediated degradation is dependent on Cullin-Ring E3 Ligases (CRLs). In order to show E3 dependency, we used the small molecules targeting VHL (VH-298^21^) as well as CRBN (Lenalidomide) as competitor molecules for E3 ligase binding. Inhibition of deneddylation conveyed by the Constitutive Photomorphogenesis 9 Signalosome (CSN/COPS9) complex subunit CSN5 can be inhibited using CSN5i-3, leading to an increase in neddylated CRLs and would be expected to increase CRL activity^22^. Polyubiquitinated targets can either bind directly to the proteasome or undergo prior unfolding via VCP/p97. To decipher the role of a VCP/p97-dependend unfolding event, the VCP/p97 inhibitor NMS-873 was used^23^. Finally, diverse proteasome inhibitors were tested and compared. While RA190 was published to covalently modify Rpn13, a polyubiquitin recruiting domain of the proteasome, MG-132, Oprozomib, Bortezomib and Ixazomib inhibit the final degradation step of the PROTAC mediated UPS pathway. Here, we grouped the proteasome inhibitors into group a) peptide-based inhibitors and b) boronic acid-containing inhibitors. Using this chemical toolbox, we determined the functional activity of the UPS pathway inhibitors to optimise the use of these inhibitors in providing information about the functional molecular mechanism of action of the underlying CRBN and VHL degrader molecules, the best concentration for using them as determined by EC_50_ values, as well as their effect on cell health. Therefore, utilization of this system can help to adjust the pathway inhibitor concentrations to the respective target system to ensure the least artificially induced side-effects possibly causing artefacts during degrader SAR studies.

## Limitations

For this study we decided to use a well-established, endogenously HiBiT-tagged cell line to study BRD4 abundance in a cellular context. The generated data are therefore dependent on the utilized cell line and the chosen target as well as utilized degradation machinery. Although our findings were transferrable from CRBN-dependent to a VHL-dependent degradation system, and from a VHL-dependent to a CRBN-dependent system, this transferability cannot be assumed for novel E3 ligase ligands which e.g. differ in their expression level, efficiency or mechanism (e.g. BIRC proteins which undergo auto degradation upon utilization^24^). However, this study demonstrated the utility of a number of tool compounds when used within a non-toxic concentration range and provides guidelines how to individually determine the cellular EC_50_s of these compounds in every system based on the presented system.

## Results

To evaluate the 12 selected inhibitors (Figure 1) that interfere with different key steps of POI degradation of the ubiquitin-based degradation pathway, we used a well-established assay system^25^. The system is based on endogenously HiBiT-tagged BRD4 in human embryonic kidney cells (HEK293T^BRD4-HiBiT^)^17^. To study the global influence of UPS inhibitors, this HiBiT system was used to measure BRD4 protein abundance in live as well as in lysed cells. Our assay design was based on BRD4 degradation induction followed by subsequent rescue of POI degradation mediated by UPS inhibitor treatment. To induce BRD4 degradation, two well-established PROTACs for BRD4, MZ1 and dBET6 were used. Using the HIBIT-tagged BRD4 system allowed us to monitor the effect of UPS inhibitors on key parameters typically used to characterise the efficiency of a PROTAC such as the concentration required for half maximum (DC_50_) and complete degradation (DC_100_) with a highly sensitive luminescence-based reporter. To monitor the time dependence of these parameters, we performed the experiment at five time points as shown in Figure 2 A. As expected, both PROTACs revealed high degradation potencies with a maximum degradation efficiency (D_max_) of nearly 100 % and DC_50_ of 20 – 50 nM for both degraders at time points from 2 – 8 h. MZ1 showed a drastic increase in potency after 24 h with a DC_50_ of approximately 30 pM using this assay system. In agreement with the literature, a “hook effect” (decrease of PROTAC efficiency at high concentrations due to preferred binary complexes^26^) was observed for dBET6, but not for MZ1. The DC_100_ was chosen as the lowest concentration at which complete degradation was observed at the examined time points. The DC_100_ values were 500 nM for dBET6 and 1 μM for MZ1, respectively. These concentrations were chosen for subsequent rescue experiments using the selected 12 UPS inhibitors. The EC_50_ values of the UPS inhibitors were determined at 4 different time points, allowing to monitor kinetic aspects of PROTAC-induced BRD4 degradation at different stages of this pathway. Data measured at 2 h are shown in Figure 2 B-G and the observed BRD4 degradation is listed in Table 1. Additional data are shown in SI Figure 1 and SI Table 1-3.

**Figure 2:**
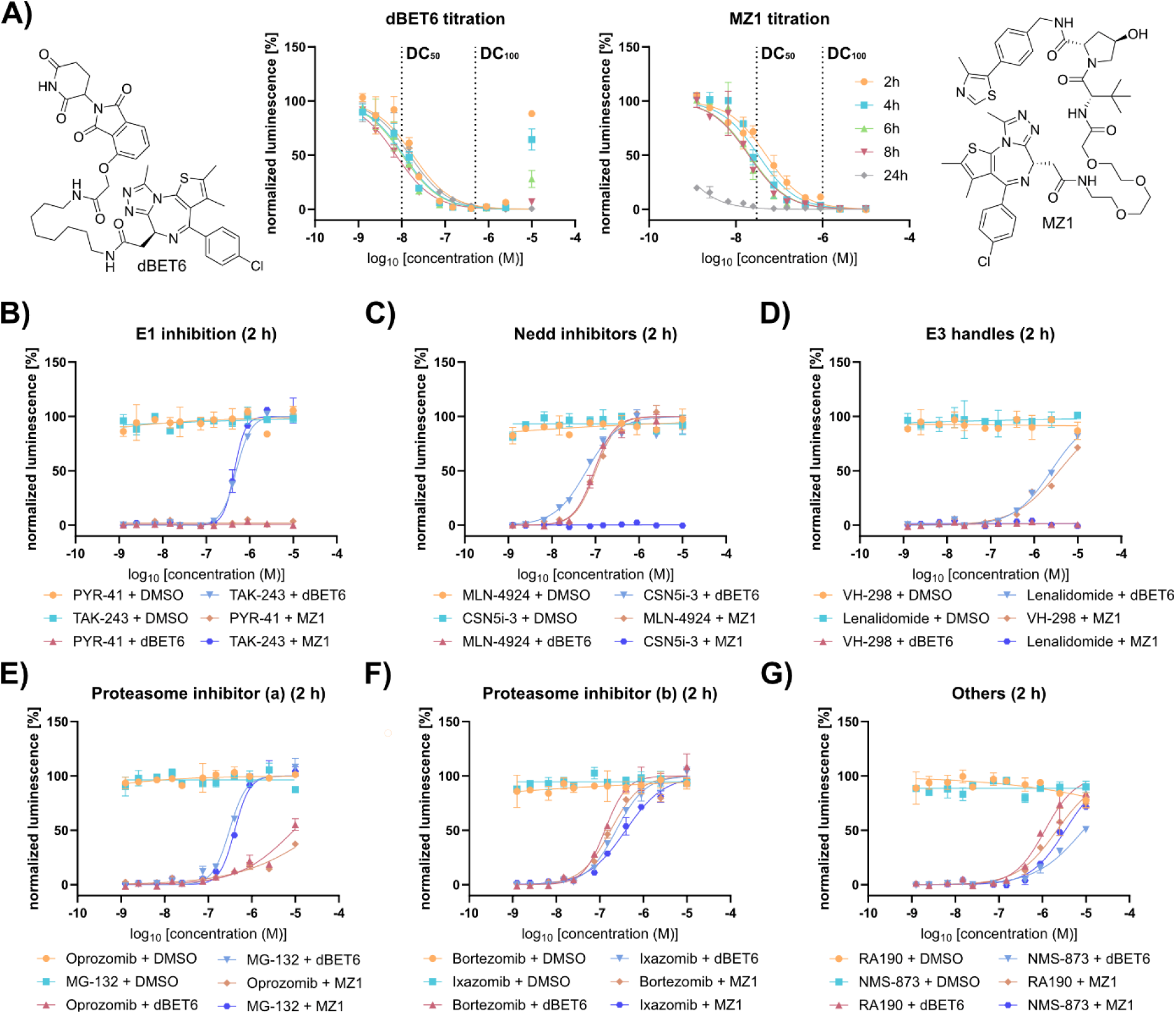
Cellular degradation (**A**) and rescue experiments (**B-G**) of BRD4 in BRD4^HiBiT^ cells. **A)** Structures and degradation curves of dBET6 (left) and MZ1 (right). Dotted lines display the concentration of half degradation (DC_50_) and maximum degradation (DC_100_). Data were measured in biological duplicates for 2, 4, 6, 8 and 24 h with error bars depicting the SD. (n=2). **B)** Rescue experiments using the E1 inhibitors TAK-243 and PYR-41. **C)** Rescue experiments using the neddylation inhibitor MLN-4924 and deneddylation inhibitor CSN5i-3. **D)** Rescue experiments using the competing E3 ligase binder for PROTAC competition with Lenalidomide for dBET6 and VH-298 for MZ1. **E)** Rescue experiments using the peptide-based proteasome inhibitors MG-132 and Oprozomib (Proteasome inhibitors group a). **F)** Rescue experiments using the boron-containing proteasome inhibitors Bortezomib and Ixazomib (Proteasome inhibitors group b). **G)** Rescue experiments using the Rpn13 inhibitor RA190 and the VCP/p97 inhibitor NMS-873. All rescue experiments were carried out with and without PROTAC co-treatment and are depicted as measurements of biological duplicates expressing the SD. (n=2). Complete data set is depicted in SI Figure 1.

**Table 1:** Results after 2h of measuring the BRD4 rescue after PROTAC and inhibitor treatment. EC50 values display the concentration at which BRD4 degradation is 50 % rescued, while the maximum recovery displays the maximum percentage of BRD4 which could be rescued using the different inhibitors. Data was extracted from three parameter fits of biological duplicates. (n=2) Results for all time points are depicted in the supplementary tables 1-3.

| 2 h | + 500 nM dBET6 | | + 1 μM MZ1 | |
| --- | --- | --- | --- | --- |
|  | EC<sub>50</sub> [nM] | Max. recovery [%] | EC<sub>50</sub> [nM] | Max. recovery [%] |
| Bortezomib | 130.9 | 100 | 177.5 | 100 |
| Oprozomib | 9311 | 55 | >25000 | 38 |
| Ixazomib | 237.8 | 100 | 393.5 | 93 |
| MG-132 | 307.9 | 100 | 406.1 | 100 |
| PYR-41 | - | 0 | - | 0 |
| TAK-243 | 491.2 | 100 | 445.3 | 100 |
| VH-298 | - | 0 | 3764 | 71 |
| Lenalidomide | 2469 | 81 | - | 0 |
| RA190 | 1144 | 84 | 2154 | 75 |
| MLN-4924 | 94.0 | 100 | 101.6 | 100 |
| CSN5i-3 | 62.3 | 100 | - | 0 |
| NMS-873 | 8712 | 50 | 3333 | 72 |
| DMSO | - | 0 | - | 0 |

**Table 2:** Recommended UPS inhibitors and doses for different time points to obtain degradation inhibition of > 75 %.

|  | 2 h | 4 h | 8 h | 24 h |
| --- | --- | --- | --- | --- |
| Bortezomib | 250 nM | 250 nM | 250 nM | 500 nM* |
| MG-132 | 750 nM | 750 nM | 750 nM | 500 nM* |
| TAK-243 | 750 nM | 750 nM | 750 nM | 150 nM* |
| VH-298 (VHL only) | 10 μM | 10 μM | 10 μM | - |
| Lenalidomide (CRBN only) | 10 μM | 10 μM | 10 μM | 10 μM |
| MLN-4924 (CRL only) | 250 nM | 250 nM | 250 nM | 250 nM |
| CSN5i-3 (Cul4 only) | 750 nM | 750 nM | 750 nM | 750 nM |

Our data showed that inhibition of E1 enzymes by TAK-243 (MLN7243) and thus initiation inhibition of the UPS, completely abrogated PROTAC-induced BRD4 degradation, resulting in complete rescue of BRD4 levels. The apparent EC_50_ values of TAK-243 were ∼ 500 nM. Furthermore, TAK-243 repressed the PROTAC-mediated BRD4 degradation over 8 h without change of potency (Table 1 and SI Table 1-2). Surprisingly however, the second E1 inhibitor PYR-41 did not abrogate PROTAC-mediated BRD4 degradation at any time point. After 24 h treatment, maximum BRD4 degradation decreased to 40 %. This effect mirrored degradation levels in TAK-243 treated cells in the absence of BRD4 targeting PROTACs (DMSO control), suggesting unspecific BRD4 degradation, possibly caused by compound toxicity at late time points (SI Figure 1 A and SI Table 3).

Next, we tested inhibitors such as MLN-4924, interfering with neddylation, a critical activation step of the recruited E3 ligases^27^. The modification of cullin by NEDD8 conjugation is specifically required for the activation of cullin-dependent E3 ligases, and potent neddylation inhibitors that have been developed are therefore valuable chemical tools to demonstrate the dependence on a cullin E3 system such as CRBN or VHL, required for the two selected BRD4 PROTACs. In contrast, deneddylation, which can be inhibited by the inhibitor CSN5i-3, is expected to result in constitutively active cullin E3 ligases and thus increase the efficacy of PROTACs that utilize cullin E3-ligases. As expected, inhibition of neddylation leads to potent inhibition of CRBN and VHL-mediated degradation activity (Figure 2 C). Interestingly, upon inhibition of the deneddylase subunit of the COPS9 complex using CSN5i-3, no effect on BRD4 degradation was observed for MZ1, a VHL recruiting PROTAC. In contrast, the most potent inhibition of BRD4-targeing degradation of all inhibitors studied was observed for the CRBN-dependent dBET6 with an EC_50_ of 60 nM (Table 1). Due to dependence of VHL on neddylation as shown by MLN-4924 treatment, resistance to CSN5 inhibition indicated a different mechanism of deneddylation. While treatment at 2 h-8 h provided reliable data on PROTAC-mediated BRD4 degradation, treatment at 24 h resulted in increased BRD4 levels which was also observed for the DMSO controls in the absence of PROTACs highlighting degradation of BRD4 by other mechanisms (SI Figure 1 B).

As a strategy demonstrating target engagement with the E3 substrate binding site, we used the monovalent E3 ligase handles (VH-298 and Lenalidomide) as competitors to compete with PROTAC binding therefore inhibiting the degradation efficiency. As expected, Lenalidomide exclusively rescued dBET6-mediated degradation while VH-298 solely rescued MZ1-mediated BRD4 degradation (Figure 2 D). Interestingly, the measured EC_50_ values were weaker than expected based on binding data measured by competition with BRET probes. This highlights the impact the catalytic turnover of PROTACs outcompeting binary complex formation (SI Figure 1 C and SI Table 1-3).

The last step in the UPS cascade is the degradation of the POI by the proteasome. To probe this step, we selected two major classes of proteasome inhibitors. Peptide-based inhibitors (Group a)) including Oprozomib and the tool compound MG-132 (Figure 2 E) which were tested along with the boronic acid-containing covalent inhibitors (Group b)) such as Bortezomib and Ixazomib (Figure 2 F). Interestingly, MG-132 displayed much higher potency with EC_50_ values around 400 nM compared to Oprozomib which showed EC_50_ values > 5 μM (Table 1 and SI Table 1-3). MG-132 treatment alone, in the absence of BRD4 targeting PROTAC caused unspecific BRD4 downregulation at 8 h and 24 h, most likely due to unspecific effects, rendering both compounds, MG-132 and Oprozomib as unsuitable control compounds for BRD4-PROTAC validation when used at concentrations >1 μM in our setup (SI Figure 1 D). Boron-containing proteasome inhibitors showed notably higher potency (EC_50_ < 300 nM) (Table 1 and SI Table 1-3). Additionally, an incubation time of 8 h did not reveal any unspecific downregulation of BRD4 protein levels while treatment for 24 h caused BRD4 downregulation in a comparable manner to MG-132 treatment (SI Figure 1 E). To our surprise, all proteasome inhibitors did not completely rescue PROTAC mediated BRD4 degradation after 24 h due to a general decrease in POI protein levels at higher proteasome inhibitor concentrations (SI Table 3). In addition to the investigated proteasome inhibitors, a VCP/p97^23^ and a covalent Rpn13^28^ inhibitor was tested (Figure 2 G). Since NMS-873 has been annotated as VCP/p97 inhibitor, we hypothesised that inhibition of the complex could reveal a protein unfolding step prior to proteasomal degradation. Indeed, treatment with NMS-873 successfully rescued BRD4 degradation, supporting this hypothesis and demonstrated the suggested role of this compound in the UPS (SI Figure 1 F). RA190 has been described in the literature as covalent Rpn13 modifier, a proteasomal subunit responsible for polyubiquitin recruitment^28^. As expected, due to the presence of a highly reactive covalent warhead, off-target activity was observed, as evidenced by the PROTAC-independent downregulation of BRD4 levels after only 4 hours of incubation. These non-specific off-target effects have recently also been reported by Dickson *et al*. (SI Figure 1 F)^29^.

Surprisingly, inhibition by E1 enzymes or proteasomal inhibition for 24 h induced unspecific BRD4 downregulation or a decrease in the capacity of the selected inhibitors to rescue POI degradation. To study this effect in more detail, we measured the kinetics of BRD4 degradation in live cells over a time course of 30 h. Remarkably, nearly all pathway inhibitors that were active in our rescue experiments significantly downregulated BRD4 in the absence of PROTACs as shown in SI Figure 2. The only inhibitors which did not non-specifically influence BRD4 levels were PYR-41, shown to be inactive in rescue experiments and both E3 ligase handles VH-298 and Lenalidomide (Figure 3 A and SI Figure 2). Inhibition of CRL neddylation by treatment with MLN-4924 increased target protein levels while the lowest concentration of 0.1 μM led to the largest increase of cellular BRD4 levels (∼40 %). Since this compound is frequently used in rescue experiments to demonstrate a CRL-dependent mechanism, this increase might lead to false positive rescue data that are most likely caused by the accumulation of BRD4 due to inhibition of its endogenous protein turnover. However, the effect may be specific to certain proteins such as BRD4 which is highly regulated by CRL whereas targets regulated by other mechanisms than CRLs may not be affected (SI Figure 2). Treatment with CSN5i-3 and therefore inhibition of deneddylation rapidly decreased total BRD4 levels up to 60 % after 8-10 h without further depletion at later time points (SI Figure 2). Inhibition of E1 enzymes resulted in the strongest unspecific modulation of cellular BRD4 protein abundance, causing rapid and complete depletion of this POI after 24 hours at TAK-243 concentrations of 10 μM and 1 μM, respectively. However, at inhibitor concentrations of 0.1 μM, BRD4 levels were downregulated by only 50 % after 24h, therefore showing a similar behaviour as seen for the studied proteasome inhibitors (SI Figure 2).

**Figure 3:**
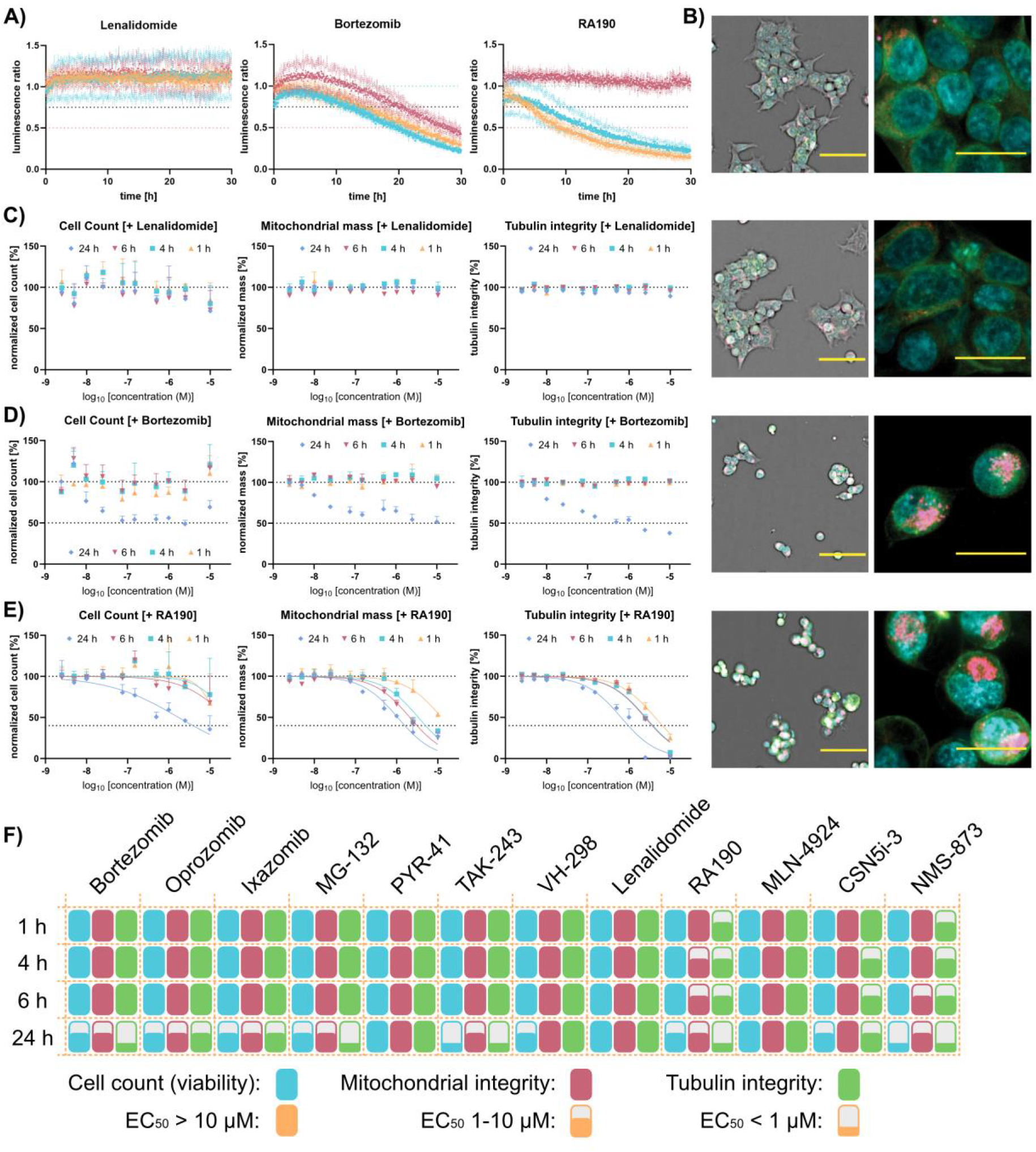
Cellular degradation kinetics (**A**) and multiplex high-content imaging data. **A)** Live cell kinetic data of HEK293T^BRD4-HiBiT^ cells stably expressing the LgBiT protein. Data were measured continuously for 30 h and collected in biological triplicates (n = 3) with dotted lines depicting the SD. **B)** The left side shows brightfield confocal images at 10X of stained (blue: DNA/nuclei, green: microtubule, red: mitochondria, and magenta: Annexin V apoptosis marker) HEK293T cells after 24h. Scale bar indicates 100 μm. Shown on the right are confocal images at 60X maginification of the same cells. Scale bar indicates 25 μm. The first images are cells treated with 0.1 % DMSO as a control compound, followed by Lenalidomid [1 μM], Bortezomib [1 μM] and RA190 [1 μM]. **C)** Normalized cell count (%) of HEK293T cells exposed to Lenalidomid, Bortezomib and RA190 at ten different concentrations ranging from 10 μM to 2.5 nM over different time-points (1h, 4h, 6h and 24h). Data was normalized against 0.1 % DMSO control. Error bars show standard error of mean (SEM) of biological duplicates. **D)** Normalized influence on mitochondrial mass (%) of HEK293T cells exposed to Lenalidomid, Bortezomib and RA190 at ten different concentrations ranging from 10 μM to 2.5 nM over different time-points (1h, 4h, 6h and 24h). Data was normalized against 0.1 % DMSO control. Error bars show standard error of mean (SEM) of biological duplicates. **E)** Normalized effect on tubulin integrity (%) of HEK293T cells exposed to Lenalidomid, Bortezomib and RA190 at ten different concentrations ranging from 10 μM to 2.5 nM over different time-points (1h, 4h, 6h and 24h). Data was normalized against 0.1 % DMSO control. Error bars show standard error of mean (SEM) of biological duplicates. **F)** Normalized cell health effects (cell count, mitochondrial integrity, tubulin integrity) of HEK293T cells exposed to all reference compounds at ten different concentrations ranging from 10 μM to 2.5 nM over different time-points (1h, 4h, 6h and 24h). The degree of filling indicates the EC_50_ values. Data was normalized against 0.1 % DMSO control.

Peptide-based proteasome inhibitors, MG-132 and Oprozomib, displayed dose-response curves with comparably small unspecific effects on cellular BRD4 levels at 0.1 μM inhibitor concentration in contrast to 1 μM and 10 μM, where drastic unspecific target regulation was observed. Interestingly, the boronic acid derivatives did not display dose response effects (Figure 3 A). Here, treatment with 0.1 μM led to similar target depletion as 10 μM, indicating a maximum inhibition of the proteasome which is also observed for treatment with 10 μM MG-132 and Oprozomib. Lastly, RA190 and NMS-873 decreased cellular BRD4 levels when used at 1 μM and 10 μM, respectively while treatment at 0.1 μM showed no influence on BRD4 protein levels (Figure 3 A).

Since the unspecific regulation of POI protein levels may be influenced by compound toxicity, we performed a high-content multiplex imaging assay to monitor the effects of the used inhibitors on cellular health as indicated by the induction of cell death pathways and the integrity or morphology of mitochondria, the nucleus and the cytoskeleton (Figure 3 B)^30^. To assess these effects, the inhibitors were tested in a dose-dependent manner in the absence of PROTACs at different time points (1 h, 4 h, 6 h and 24 h). First, auto fluorescence or compound precipitation was assessed detecting Hoechst high intensity objects^31^. Here, none of the used inhibitors showed detectable autofluorescence or precipitation at the concentrations used (SI Table 4). Next, cellular viability was assessed based on the nuclear morphology followed by testing for alterations of tubulin structure and potential increase of mitochondrial mass. Within this compound set, three major general effects on cellular health were detected: First, Lenalidomide showed no effect on cell viability, mitochondrial mass and tubulin integrity (Figure 3 C). Second, inhibitors that displayed general toxicity and additionally induced changes in mitochondrial mass and tubulin integrity as seen for example for Bortezomib (Figure 3 D and SI Figure 3). Lastly, compounds that significantly altered mitochondrial mass or tubulin integrity without affecting the cell count or inducing toxicity.

In order to evaluate the effect of the inhibitors on the used cell system, we investigated their effect in a concentration- and time-dependent manner. These studies may provide insights into unspecific BRD4 downregulation which may be caused by effects on cell viability. In this screen, NMS-873 was found to significantly disrupt tubulin integrity at concentrations and time points where no effect on the general viability was observed. An increase of mitochondrial mass was also observed at earlier time points than a decrease of the normalized cell count, indicating that the impact on general health of this compound could be attributed to an interaction of the molecule on microtubules. Nevertheless, general viability and disruption or binding of cell organelles are a complex interplay of diverse factors, which makes it difficult to provide absolute statements. For RA190, we observed a rapid increase of mitochondrial mass and a decrease in tubulin integrity before affecting the overall cell count in a comparable manner (Figure 3 E). The rapid impact of this compound on the integrity of these cellular structures, ruled out that these effects are caused by underlying general toxicity but suggested direct modulation of mitochondria and tubulin function (SI Figure 3 B and C). Additionally, later time points show increasing and dose-dependent effects on the mitochondrial mass and tubulin disruption supporting this hypothesis (SI Figure 3). Generally, we found that none of the inhibitors had a significant effect on cell count (EC_50_ < 10 μM) at early time points (1 h to 6 h) while after 24 h, nearly all inhibitors affected cell viability (as generally observed in the multiplexed and HiBiT assay), possibly explaining the unspecific alteration of BRD4 abundance upon treatment. This is most likely due to the drastic increase in cellular debris caused by the inhibited proteasomal machinery (Figure 3 F). Changes in tubulin integrity or increasing mitochondrial mass at significantly lower doses of compounds and early applications, can cause an influence on cell viability at later time points, as shown by MG-132, RA190 and NMS-873 (Figure 3 F). Therefore, the use of these compounds should be avoided as UPS control compounds (in case of RA190 and NMS-873) or low concentrations, preferably with additional kinetic studies should be used.

Using the combined data, we compiled recommendations for selecting the most suitable pathway inhibitors together with their appropriate concentrations and time points for our CRBN and VHL-mediated BRD4 degradation system. In this comparative study we considered a rescue as ‘significant’, if the residual degradation at DC_50_ concentration was lower than 10 %. We chose MZ1 and dBET6 as prototypes of an extremely potent PROTACs and added dBET1 (Figure 4 A) as additional test degrader, due to its reduced degradation efficiency to validate our recommended concentrations also for less efficient PROTACs (Figure 4 B and C). Since to date most degraders rely on proteasomal degradation, inhibitors for the proteasome mark an important tool for degrader evaluation. Bortezomib was found to be the most potent proteasome inhibitor of PROTAC-induced protein degradation with an EC_50_ of roughly 130 nM (Table 1). We found that the frequently chosen concentration of 1 μM already exhibited off-pathway effects expressed in a decrease of cell viability, we therefore propose to lower the recommended concentration to 250 nM as at this concentration already a significant decrease in degradation was observed (Figure 4 B and C and SI Figure 4). Moreover, the use of our recommended concentration only decreased BRD4 levels after 24 h from approx. 50 % seen for 1 μM (Figure 3 A and D) to only 10 %. MG-132, a control frequently used in literature showed high concentration dependency for unspecific lowering of target protein levels but also higher EC_50_ values in the rescue experiments of around 300 nM (Table 1). For this pathway inhibitor, the weaker EC_50_ would suggest a higher recommended concentration. We found a concentration of 750 nM most optimal resulting in a comparable BRD4 rescue from PROTAC-mediated degradation as 250 nM Bortezomib, reaching the < 10 % threshold at DC_50_ (Figure 4 B and C and SI Figure 4). For TAK-243 a recommend a concentration of 750 nM was suitable in our test systems (Figure 4 C). Increase of TAK-243 concentration significantly increased the off-pathway activity and should therefore be avoided (SI Figure 2). Both E3 ligase handles VH-298 and Lenalidomide were found to be not generally toxic but they also resulted in relatively weak EC_50_ values (Table 1). Due to this property, only treatment with 10 μM of both ligands led to a decrease in PROTAC-induced degradation which matched our defined guideline. Using the CRBN and VHL E3 handles as inhibitors is therefore only recommended for target ligase validation and exclusion of CRBN/VHL independent degradation. Interestingly, treatment with 10 μM VH-298 in CRBN recruiting systems led to an increase in degradation efficiency after 24 h. Last, we compiled recommended concentrations for the inhibition of the NEDD8 cascade using MLN-4924 and CSN5i-3. Here, we chose 250 nM for MLN-4924 due to its high potency, leading to excellent abrogation of PROTAC induced degradation even after 24 h (Figure 4 C). As also seen for other assays utilizing MLN-4924 (SI Figure 1 and 2), we observed an increase in BRD4 levels upon treatment with MLN-4924 of approx. 30 %, possibly caused by modulation of an endogenous regulator of BRD4. For CSN5i-3, a concentration of 750 nM was required for a significant inhibition of PROTAC dependent degradation which was deemed to be low enough to avoid unspecific reduction of BRD4 levels but sufficient for rescuing PROTAC dependent degradation. Remarkable however, (Figure 2 C) only CRBN-dependent degradation was inhibited by CSN5i-3 while VHL-dependent degradation (MZ1) was not found to be inhibited (Figure 4 and SI Figure 4). This was unexpected due to the association of the COP9 signalosome with CUL2 ligases^32^.

**Figure 4:**
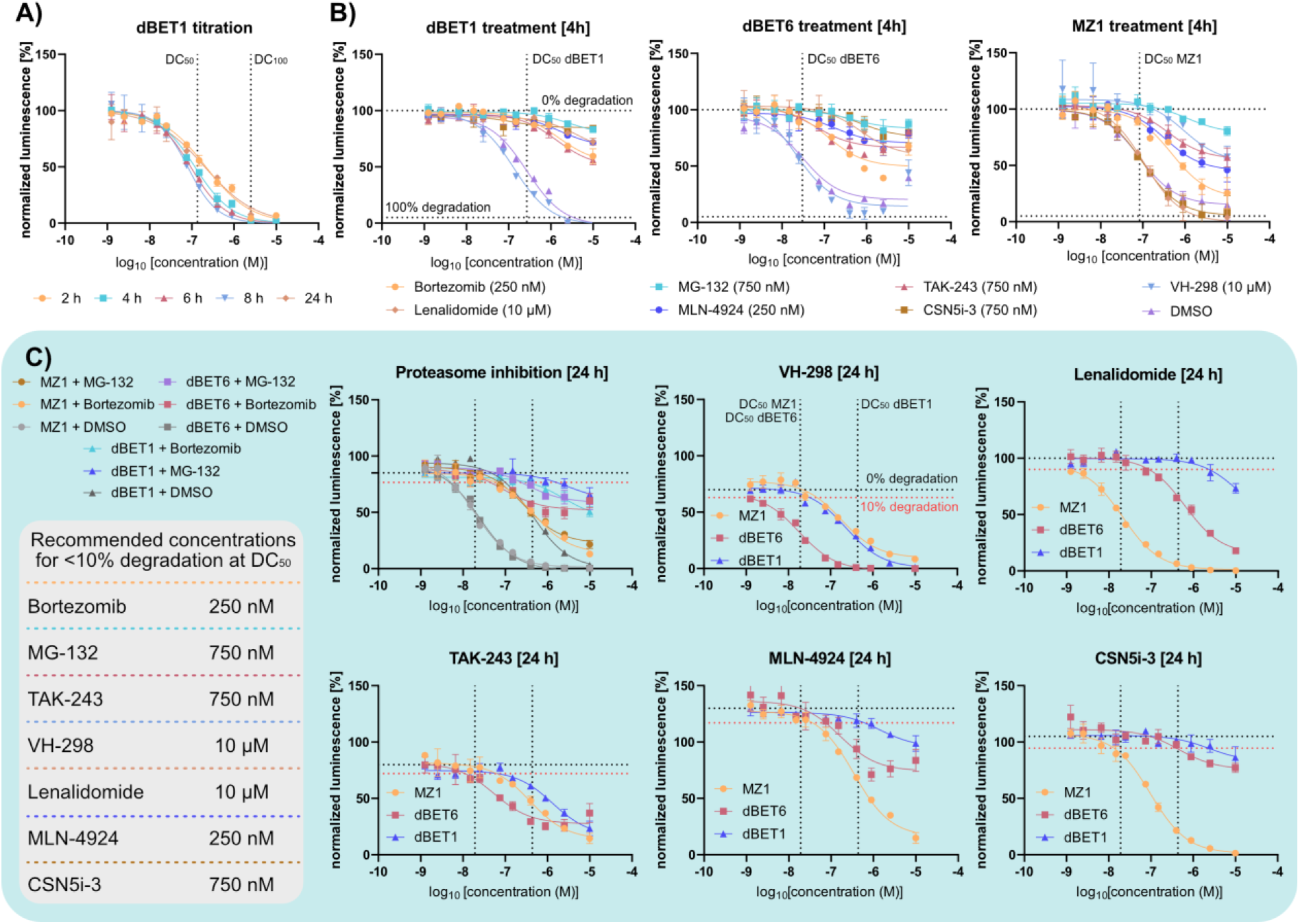
Degrader titrations after inhibitor treatment at the recommended concentrations. A) dBET1 titration to BRD4^HiBiT^ cells for DC_50_ determination at different time points. Data were measured in biological replicates with error bars expressing the SD. (n = 2). B) Degrader titrations after treatment with fixed recommended inhibitor concentration shows significant degradation reduction at the DC_50_ (vertical dotted line) after 4 h. Horizontal dotted lines indicate 0 and 100 % degradation, respectively. Data were measured in biological replicates with error bars expressing the SD. (n = 2). C) Degrader titrations after inhibitor addition at recommended concentration (grey box), measured after 24 h. Legends for the dotted lines is shown within the panel for VH-298 where the red dotted line represents the cut off for < 10 % degradation. Successful degradation rescue was achieved through < 10 % target degradation at DC_50_ concentration of the respective PROTAC (above the red dotted line at the corresponding vertical dotted line). Data were measured in biological triplicates with error bars expressing the SD. (n = 3). Complete data set is shown in SI Figure 4.

## Discussion

Inhibition of PROTAC-induced target degradation by available small molecules is now considered an essential control during the development of degrader molecules. However, the lack of data regarding the optimal inhibitor concentration - sufficient for the degrader induced inhibition but low enough to avoid toxicity – may have confounded experimental results. Here, we successfully established a functional assay system to profile different UPS inhibitors and additionally analysed these for their off-target activity within living cells. During our study, we were able to prove that PYR-41 has no effect on the UPS pathway, therefore being a false-positive in the literature^18^. Additionally, TAK-243 was indeed found to be a UPS-active compound while showing very high off-target activity after more than 8 h making this control compound not suitable for studies over 24 h. The drastic off-target effects of RA190 indicates far more cellular targets of this compound as initially published which is in line with recently published findings^29^. Last, the VCP/p97 inhibitor NMS-873 was also found to have severe cellular side effects, excluding this as a possible control compound for degrader MOA determination.

Interestingly, we found that concentrations which are regularly used in the literature are already within the toxic range which might lead to unwanted side effects and assay artefacts. We therefore compiled recommended concentrations which display significantly lower working concentrations for our test case, drastically decreasing possible off-target effects, based on our newly postulated guideline for degradation rescue. Moreover, similar effects on mitochondria^33,34^ and tubulin^35^ were already reported for proteasome inhibitors, showing that the findings presented here is in agreement with available reports.

Due to the high dependence on both, target protein and E3 ligase, we generally recommend testing for unspecific target regulation upon treatment with the pathway inhibitors during the development of novel degraders. This can either be done by using this functional system or by including controls during protein abundance measurements which were treated with control compounds but without the degrader to estimate the impact on the target. Since we focused on the highly studied and robust BRD4-targeting system using the established CRBN/VHL E3 ligases, the recommended concentrations most likely differ for every new target depending on the cellular turnover and expression levels and therefore need to be adjusted to each case.

This study successfully proved that these control compounds interfere with a complex mechanism and should not be used in high concentrations without excluding unspecific target regulation. We hope to raise awareness to these unspecific effects through this study, supporting a clean validation of PROTACs and other degraders, minimizing assay artefacts.

### Significance

PROTACs are of increasing interest especially for the pharmaceutical industry. Due to the degrader molecules now entering clinical trials, thorough evaluation of these molecules is required for a complete understanding of their mechanism of action. For this, inhibitors of the ubiquitin-proteasomal system (UPS) are often used for the demonstration of E3 recruitment and UPS-dependent protein degradation and therefore exclude e.g. toxicity related effects on target protein levels. This study uses a functional assay system for the evaluation of available inhibitors which are often used in literature and provides recommended concentrations for maximum effect while maintaining the least invasive concentrations to minimize misinterpretation of possible off-target effects during degrader screening and evaluation campaigns.

### Star methods

## RESOURCE AVAILABILITY

Lead contact: Martin P. Schwalm.

## Materials availability

Inhibitors were purchased from Sigma Aldrich, Tocris or MedChemExpress and HEK293T^BRD4-HiBiT^ cells were obtained as a kind gift from Promega Corp. MZ1 was obtained from opnMe.

## Data and code availability

- Data reported in this paper will be shared by the lead contact upon request.
- This paper does not report original code.
- Any additional information required to reanalyse the data reported in this paper is available from the lead contact upon request.

## EXPERIMENTAL MODEL AND SUBJECT DETAILS

### Cell lines

HEK293T^BRD4-HiBiT^ (female, fetus) cells were regularly tested for mycoplasma infection. Cells were grown in DMEM medium supplemented with 10% FBS and 1% Penicillin/Streptomycin (100U/ml penicillin and 100 mg/ml streptomycin) at 37 C° and 5% CO_2_.

HEK293T (ATCC® CRL-1573™) (female, fetus) cells were regularly tested for mycoplasma infection. Cells were grown in DMEM plus L-Glutamine (High glucose) medium supplemented with 10% FBS and 1% Penicillin/Streptomycin (100U/ml penicillin and 100 mg/ml streptomycin) at 37 C° and 5% CO_2_.

### Inhibitors used in this study

All inhibitors were purchased from Sigma Aldrich, Tocris or MedChemExpress with fresh stocks prepared prior experiments. MZ1 was obtained from opnMe.

### Method details

#### HiBiT endpoint detection for BRD4 degradation

Endogenously BRD4 HiBiT-tagged HEK293T (HEK293T^BRD4-HiBiT^) cells were obtained as a kind gift from Promega Corp. To measure degradation, 10 μl of a total concentration of 2.5×10^5^ cells/ml in DMEM medium were seeded into white small volume 384 well plates (Greiner, 784075) and allowed to settle overnight. Subsequently, the PROTACs were titrated to the seeded cells, using an Echo acoustic dispenser (Labcyte) and the plate was incubated for the indicated time at 37°C and 5 % CO_2_. After incubation, HiBiT Lytic detection reagent was prepared by dilution of LgBiT protein (1:100) and lytic substrate (1:50) in Lytic detection buffer (Promega, N3040). For detection, 10 μl of the prepared mix was added to the treated cells and incubated for 10 minutes at room temperature. Readout was carried out in a PheraStar FSX plate reader (BMG Labtech) using the LUM plus optical module. Degradation data were then plotted with GraphPad Prism 9 software using a normalized 3-parameter curve fit with the following equation: Y = 100/(1 + 10^(X-LogIC50)).

#### Functional assay for BRD4 degradation rescue

To measure functional degradation rescue and therefore UPS inhibitors, 10 μl of HEK293T^BRD4-HiBiT^ cells with a total concentration of 2.5×10^5^ cells/ml in DMEM medium were seeded into white small volume 384 well plates (Greiner, 784075). Subsequently, UPS inhibitors were titrated to the seeded cells, using an Echo acoustic dispenser (Labcyte). Additionally, 500 nM dBET6 or 1 μM MZ1 was added to the cells for degradation induction. The plate was incubated for the indicated time at 37°C and 5 % CO_2_. After incubation, HiBiT Lytic detection reagent was prepared by dilution of LgBiT protein (1:100) and lytic substrate (1:50) in Lytic detection buffer (Promega, N3040). For detection, 10 μl of the prepared mix was added to the treated cells and incubated for 10 minutes at room temperature. Readout was carried out in a PheraStar FSX plate reader (BMG Labtech) using the LUM plus optical module. Degradation data were then plotted with GraphPad Prism 9 software using a four parameter fit with the equation Y=Bottom + (Top-Bottom)/(1+(IC50/X)^HillSlope).

#### BRD4 degradation kinetics

For cellular degradation kinetics, HEK293T^BRD4-HiBiT^cells were transfected in a T75 flask, using FuGENE 4K (PROMEGA) following the manufacturer’s protocol with the LgBiT. After allowing the cells to express the LgBiT protein for 20 h at 37 °C and 5% CO_2_, the cells were harvested and the medium was exchanged to CO_2_ independent medium (Life Sciences) containing Endurazine substrate (PROMEGA) according to the manufacturers protocol. The cells were either transferred into white small volume 384 well plates (Greiner, 784075) and incubated for 2.5h prior readout for substrate activation. After incubation, compounds were added and the plate was sealed with BreathEasy plate seal (Sigma Aldrich Z380059) and continuously measured every 5 min for 30h in a PHERAstar FSX plate reader (BMG Labtech) using the LUM plus module. Untreated baseline measurements were subtracted from the measurements to normalize the data and graphed, using GraphPad Prism 9.

#### Multiplex high-content viability assessment

To assess the compounds influence on cell health, a high-content screen in living cell called multiplex, as described previously by Tjaden et al.^31^ was performed. In brief, HEK293T (ATCC® CRL-1573™) were cultured in DMEM plus L-Glutamine (High glucose) supplemented by 10% FBS (Gibco) and Penicillin/Streptomycin (Gibco). Cells were seeded at a density of 1250 cells per well in a 384 well plate in culture medium (Cell culture microplate, PS, f-bottom, μClear®, 781091, Greiner), with a volume of 50 μL per well. All outer wells were filled with 100 μL PBS-buffer (Gibco). Simultaneously with seeding, cells were stained with 60 nM Hoechst33342, 75 nM Mitotracker red, 0.3 μL/well Annexin V Alexa Fluor 680 conjugate and 25 nL/well BioTracker™ 488 Green Microtuble Cytoskeleton dye. Cell shape and fluorescence was measured 1h, 4h, 6h and 24 hours after compound treatment using the CQ1 high-content confocal microscope (Yokogawa, Musashino, Japan). The following setup parameters were used for image acquisition: Ex 405 nm/Em 447/60 nm, 500ms, 50%; Ex 561 nm/Em 617/73 nm, 100 ms, 40%; Ex 488/Em 525/50 nm, 50 ms, 40%; Ex 640 nm/Em 685/40, 50 ms, 20%; bright field, 300 ms, 100% transmission, one centered field per well, 7 z stacks per well with 55 μm spacing. The compounds were tested at ten different concentrations ranging from 10 μM to 2.5 nM. Acquired images of the cells were processed using the Yokogawa CellPathfinder software (v3.04.02.02). Cells were detected and gated with a machine learning algorithm as described previously^30^. Results were normalized to cells exposed to 0.1 % DMSO. All Compounds were tested in biological duplicates and SEM (standard error of mean) was calculated of biological duplicates.

## Author contribution

Manuscript was prepared by MPS and edited by all authors. Study design and cellular assays were performed by MPS. Viability and toxicity assays were performed by AM. Compound management was conducted by FAG. LE conducted experiments not included in the manuscript. Scientific supervision by MBR, SMK and SK.

**Acknowledgement**

MPS, AM, FAG, LE, SM and SK are grateful for support by the Structural Genomics Consortium (SGC), a registered charity (no: 1097737) that receives funds from Bayer AG, Boehringer Ingelheim, Bristol Myers Squibb, Genentech, Genome Canada through Ontario Genomics Institute, EU/EFPIA/OICR/McGill/KTH/Diamond Innovative Medicines Initiative 2 Joint Undertaking [EUbOPEN grant 875510], Janssen, Pfizer and Takeda and by the German Cancer Research Center DKTK and the Frankfurt Cancer Institute (FCI). MPS is funded by the Deutsche Forschungsgemeinschaft (DFG, German Research Foundation), CRC1430 (Project-ID 424228829). The CQ1 microscope was funded by FUGG (INST 161/920-1 FUGG).

