## Supplemental Information for "Functional characterization of pathway inhibitors for the ubiquitin-proteasome system (UPS) as tool compounds for CRBN and VHL-mediated targeted protein degradation"

### Pathway inhibitors for CRBN and VHL-based target degradation and how to use them

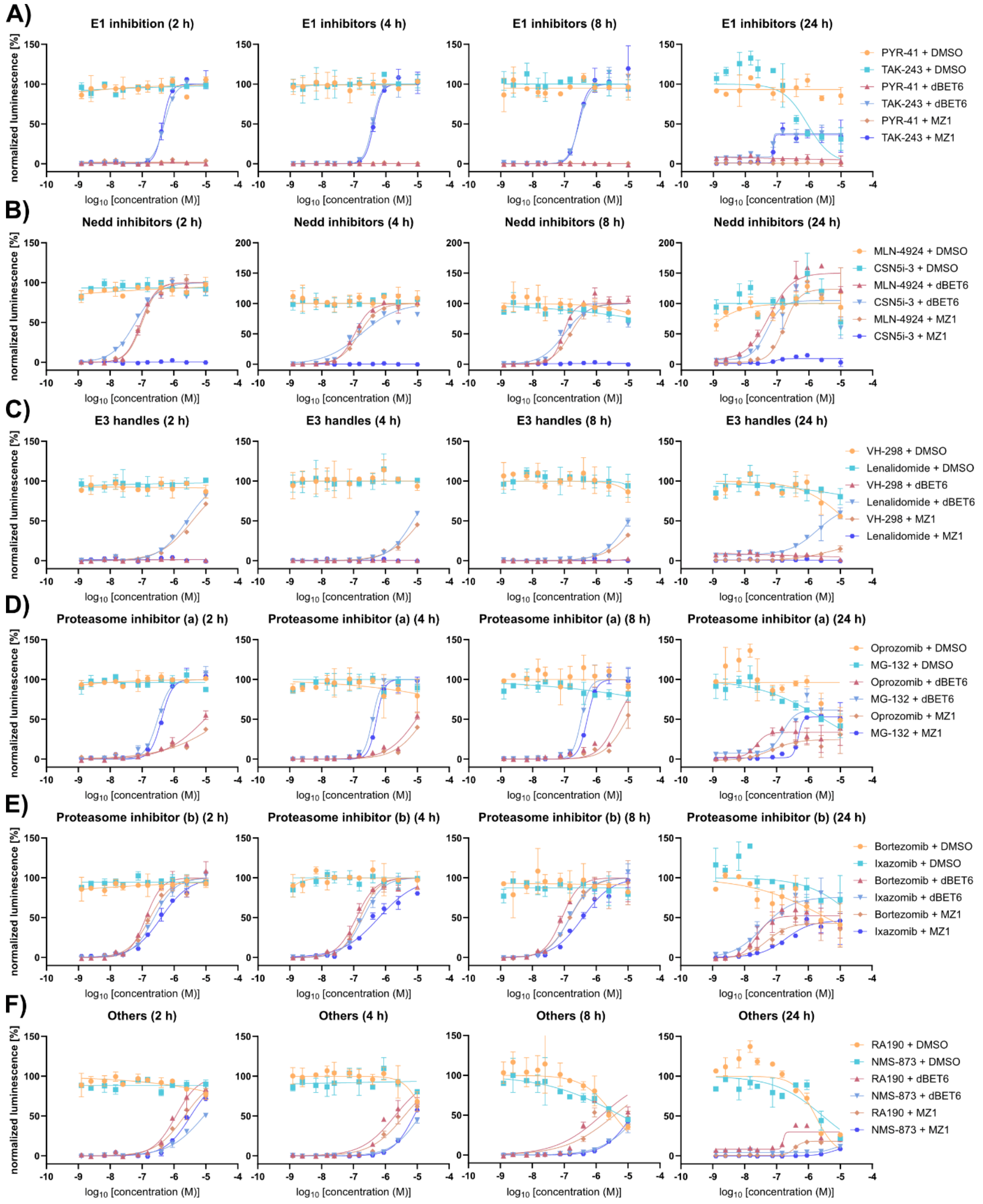


SI Figure 1: Rescue experiments of either DMSO, dBET6 or MZ1 treated BRD4$^{HiBiT}$ cells, measured after 2, 4, 8 and 24 h (left to right). **A)** Rescue experiments using the E1 inhibitors TAK-243 and PYR-41. **B)** Rescue experiments using the neddylation inhibitor MLN-4924 and deneddylation inhibitor CSN5i-3. **C)** Rescue experiments using the competing E3 ligase binder for PROTAC competition with Lenalidomide for dBET6 and VH-298 for MZ1. **D)** Rescue experiments using the peptide-based proteasome inhibitors MG-132 and Oprozomib (Proteasome inhibitors group a). **E)** Rescue experiments using the ron-containing proteasome inhibitors Bortezomib and Ixazomib (Proteasome inhibitors group b). **F)** Rescue experiments using the Rpn13 inhibitor RA190 and the VCP/p97 inhibitor NMS-873. All rescue experiments were carried out with and without PROTAC co-treatment and are depicted as measurements of biological replicates expressing the SD. (n=2)

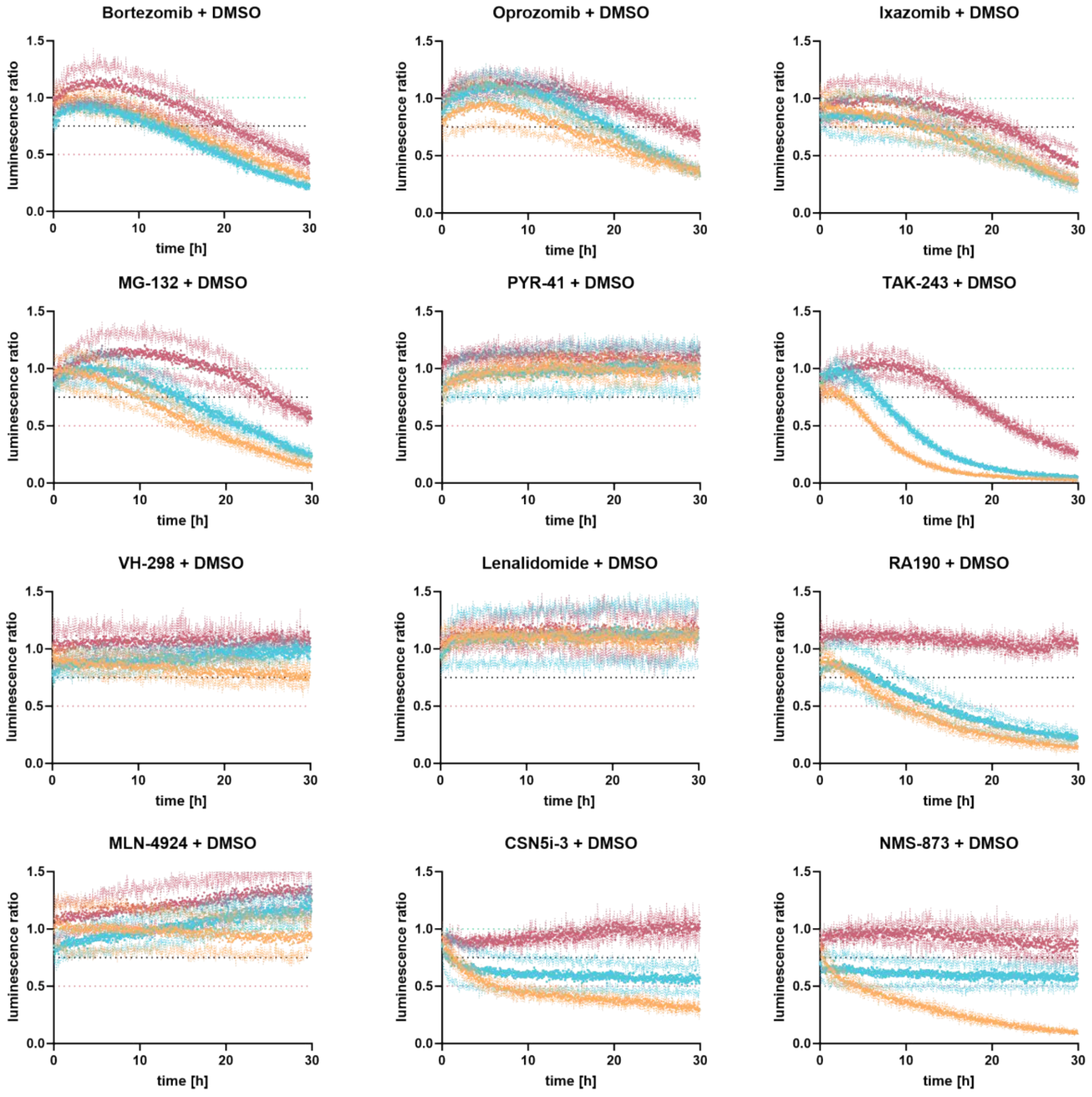


SI Figure 2: Kinetic monitoring of cellular BRD4 levels in living cells. LgBiT expressing HEK293T$^{BRD4\text{-}HiBiT}$ cells were monitored over a time course of 30 h after inhibitor treatment with 10 µM (yellow), 1 µM (blue) and 100 nM (red) of the indicated inhibitor. The dotted lines represent 100 % BRD4 levels (green), 75 % BRD4 levels (black) and 50 % BRD4 levels (red). Data were measured in biological replicates with error bars shown as dots expressing the SD. (n = 2)

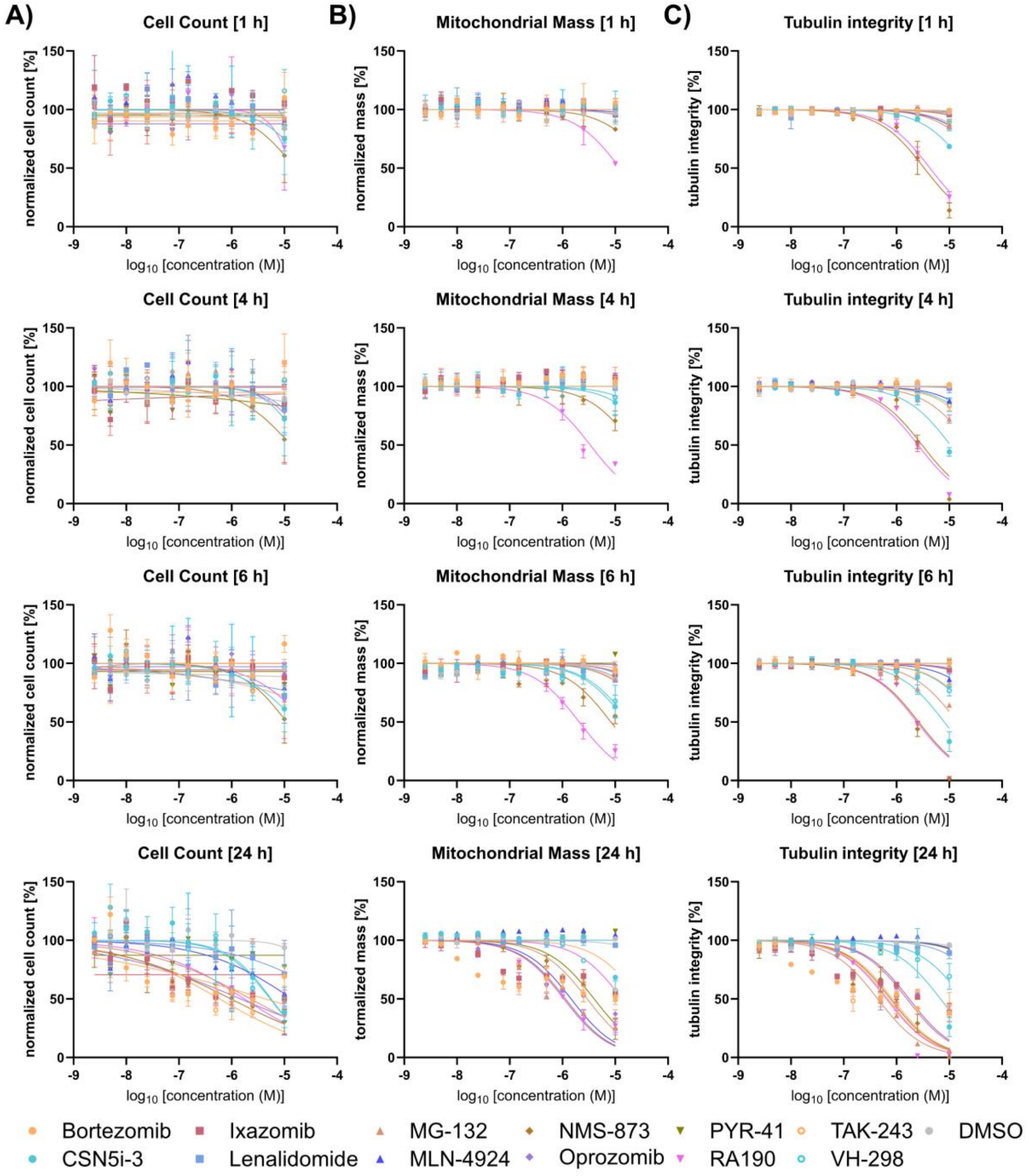


SI Figure 3: A) Normalized cell count (%) of HEK293T cells exposed to all reference compounds at ten different concentrations ranging from 10 µM to 2.5 nM over different time-points (1h, 4h, 6h and 24h). Data was normalized against 0.1 % DMSO control. Error bars show standard error of mean (SEM) of biological duplicates. B) Normalized influence on mitochondrial mass (%) of HEK293T cells exposed to all reference compounds at ten different concentrations ranging from 10 µM to 2.5 nM over different time-points (1h, 4h, 6h and 24h). Data was normalized against 0.1 % DMSO control. Error bars show standard error of mean (SEM) of biological duplicates. C) Normalized effect on tubulin integrity (%) of HEK293T cells exposed to all reference compounds at ten different concentrations ranging from 10 µM to 2.5 nM over different time-points (1h, 4h, 6h and 24h). Data was normalized against 0.1 % DMSO control. Error bars show standard error of mean (SEM) of biological duplicates.

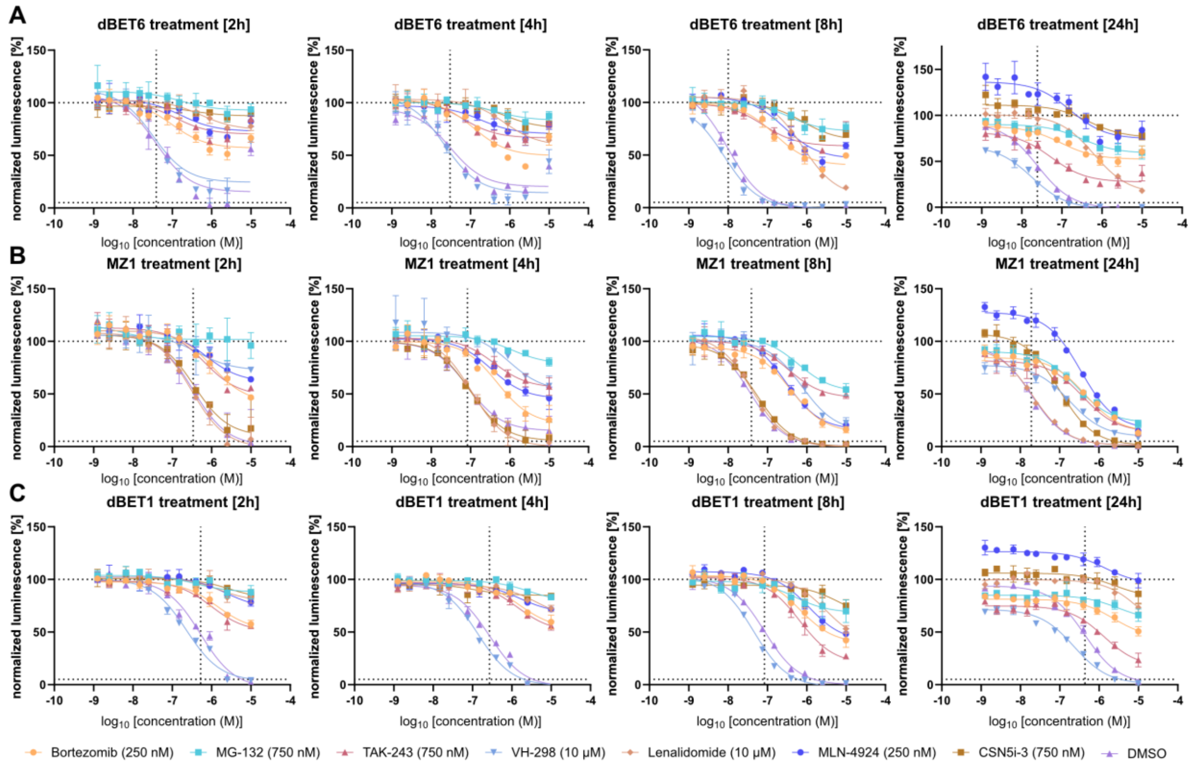


SI Figure 4: Test of recommended inhibitor concentrations using dBET6 (A), MZ1 (B) and the less effective dBET6 analogue dBET1 (C). Data were measured after 2, 4, 8 and 24 h for all combinations using the HEK293T$^{BRD4\text{-}HiBiT}$ cell line. Dotted lines indicate the DC50 (vertical) and 0/100 % protein level (horizontal). 2-8 h data were measured in biological replicates with error bars expressing the SD (n = 2), 24 h data were measured in biological triplicates with error bars expressing the SD (n = 3)

SI Table 1: Results after 4h of measuring the BRD4 rescue after PROTAC and inhibitor treatment. EC50 values display the concentration at which BRD4 degradation is 50 % rescued, while the maximum recovery displays the maximum percentage of BRD4 which could be rescued using the different inhibitors. Data was extracted from three parameter fits of biological duplicates. (n=2)

| 4 h | + 500 nM dBET6 | | + 1 µM MZ1 | |
|---|---|---|---|---|
| | EC50 [nM] | Max recovery [%] | EC50 [nM] | Max recovery [%] |
| Bortezomib | 121.2 | 100 | 160.9 | 100 |
| Oprozomib | 9376 | 54 | 16290 | 40 |
| Ixazomib | 195.9 | 100 | 617.9 | 80 |
| MG-132 | 392.5 | 100 | 522.6 | 100 |
| PYR-41 | - | 0 | - | 0 |
| TAK-243 | 379.1 | 100 | 418.5 | 100 |
| VH-298 | - | 0 | 12780 | 45 |
| Lenalidomide | 7271 | 59 | - | 0 |
| RA190 | 1976 | 67 | 3877 | 62 |
| MLN-4924 | 107.9 | 100 | 148.4 | 100 |
| CSN5i-3 | 172.8 | 83 | - | 0 |
| NMS-873 | 12160 | 45 | 7872 | 58 |
| DMSO | - | 0 | - | 0 |

SI Table 2: Results after 8h of measuring the BRD4 rescue after PROTAC and inhibitor treatment. EC50 values display the concentration at which BRD4 degradation is 50 % rescued, while the maximum recovery displays the maximum percentage of BRD4 which could be rescued using the different inhibitors. Data was extracted from three parameter fits of biological duplicates. (n=2)

| | + 500 nM dBET6 | | + 1 µM MZ1 | |
|---|---|---|---|---|
| | EC50 [nM] | Max recovery [%] | EC50 [nM] | Max recovery [%] |
| Bortezomib | 84.7 | 95* | 151.2 | 100 |
| Oprozomib | 4461 | 83 | 9029 | 55 |
| Ixazomib | 148.8 | 100 | 378.1 | 100 |
| MG-132 | 336.3 | 100 | 517.6 | 100 |
| PYR-41 | - | 0 | - | 0 |
| TAK-243 | 284.6 | 114 | 271.7 | 100 |
| VH-298 | - | 0 | 23850 | 32 |
| Lenalidomide | 10560 | 49 | - | 0 |
| RA190 | 1505 | 69* | 3741 | 53* |
| MLN-4924 | 90.8 | 100 | 150.8 | 100 |
| CSN5i-3 | 98.8 | 96* | - | 0 |
| NMS-873 | 20720 | 36 | 13440 | 42 |
| DMSO | - | 0 | - | 0 |

SI Table 3: Results after 24h of measuring the BRD4 rescue after PROTAC and inhibitor treatment. EC50 values display the concentration at which BRD4 degradation is 50 % rescued, while the maximum recovery displays the maximum percentage of BRD4 which could be rescued using the different inhibitors. Data was extracted from three parameter fits of biological duplicates. (n=2)

| | + 500 nM dBET6 | | + 1 µM MZ1 | |
|---|---|---|---|---|
| | EC50 [nM] | Max recovery [%] | EC50 [nM] | Max recovery [%] |
| Bortezomib | 23.6 | 60* | 51.7 | 49* |
| Oprozomib | 23.0 | 38* | 69.8 | 25* |
| Ixazomib | 37.3 | 80 | 193.2 | 48 |
| MG-132 | 158.4 | 80* | 474.4 | 55 |
| PYR-41 | - | 0 | - | 0 |
| TAK-243 | 74.4 | 40* | 76.7 | 43 |
| VH-298 | - | 0 | 3586 | 15 |
| Lenalidomide | 1917 | 60 | - | 0 |
| RA190 | 161.2 | 38* | 361.7 | 17 |
| MLN-4924 | 56.8 | 162* | 170.3 | 136* |
| CSN5i-3 | 48.2 | 126* | 85.9 | 14* |
| NMS-873 | 5656 | 11 | 6932 | 9 |
| DMSO | - | 0 | - | 0 |
